# An updated dataset of early SARS-CoV-2 diversity supports a wildlife market origin

**DOI:** 10.1101/2025.04.05.647275

**Authors:** Zach Hensel, Florence Débarre

## Abstract

The origin of SARS-CoV-2 has been intensely scrutinized, and epidemiological and genomic evidence has consistently pointed to Wuhan’s Huanan Seafood Wholesale Market as the epicenter of the COVID-19 pandemic. Early cases were associated with this market, and environmental sequencing placed the common ancestor of SARS-CoV-2 genomic diversity within the market. Phylogenetic analysis also suggested separate introductions of lineages A and B into the human population, a finding that can be tested with additional data. Here, we curated an expanded sequence dataset of early SARS-CoV-2 viral genomes, including newly available sequences from mid-January 2020. In this dataset, we found no additional support for previously proposed alternative progenitor sequences, or for any evolutionary intermediates between lineages A and B in the human population. Instead, we identified SARS-CoV-2 lineages that may have spread from the market, and additional samples of a sublineage of lineage A with three mutations, including one found in closely related bat coronaviruses. Although our analysis of early pandemic genomes suggests that this mutation is unlikely to characterize the immediate SARS-CoV-2 ancestor, it is more plausible than two previously proposed ancestral genomes. These findings reinforce the proposed emergence of SARS-CoV-2 from the wildlife trade at the Huanan market, demonstrating how new data continues to both solidify and clarify our understanding of how the pandemic began.

## Introduction

The origin of SARS-CoV-2 has been the subject of intense scrutiny and speculation since the virus was first identified in Wuhan, China in late 2019. The earliest reported COVID-19 cases were associated with Wuhan’s Huanan Seafood Wholesale Market [1,2], immediately suggesting that SARS-CoV-2 emerged from its mammalian wildlife trade [3,4]. Subsequent evidence has consistently supported this hypothesis. Spatiotemporal epidemiological data unambiguously distinguished the area near the market as the epicenter of the outbreak [5–8]. Sequencing of environmental samples placed all known genomic diversity of SARS-CoV-2 at the end of 2019 in a small section of the market [9]. A stone’s throw away within the market, a stall was found to contain nucleic acid from both SARS-CoV-2 and animals capable of SARS-CoV-2 transmission [10–12].

Phylogenetic analysis and epidemic modeling suggested that the two clades of SARS-CoV-2 in the early pandemic (lineage A and lineage B, separated by two mutations) were founded by at least two introductions from an unsampled reservoir in proximal animal hosts [13]. Identification of samples with transitional sequences in mid-January 2020 or earlier could challenge this interpretation of the data [14]. However, most analyses to date have been limited to sequence datasets with few samples until mid-January 2020. Furthermore, the earliest samples are dominated by those collected from market-linked patients, or from a familial cluster in Guangdong province and linked to a Wuhan hospital [15].

Some have predicted that additional data could support an alternative scenario, according to which the pandemic arose from a single SARS-CoV-2 introduction carrying one additional mutation in addition to the two characterizing lineage A [16,17]. In such a scenario, the likelihood that the pandemic originated in the market is reduced, because the inferred date of the primary infection would be earlier (further from the earliest known market-linked infections), and because the proposed ancestral mutations have not been reported in market-linked sequences. Conversely, additional data could further support the market as the pandemic epicenter if mutations found in market-linked sequences are also identified in sequences without a known link to the market.

Analysis of early-pandemic sequencing data to date has largely focused on genome sequences available in the genetic sequence repository GISAID, in addition to sequences detailed in the 2021 joint WHO-China study on the origins of SARS-CoV-2 [13,18]. Motivated by the recent availability of early-pandemic sequences collected in Shanghai and Anyang, China [19,20], we searched for additional sequences not included in previously published phylogenetic analyses, including from raw sequencing data that we assembled. We included these new sequences in a curated and updated early-pandemic sequence dataset, from which we excluded sequences identified as duplicates or whose coverage was low. Importantly, new sequences in our dataset were collected in mid-January 2020, a period with few sequences in previous analyses. We did not find significant support for either of two previously proposed progenitor sequences [16,17], and we found no transitional sequences between lineage A and lineage B. Instead, we found significant support both inside and outside of Wuhan for a lineage associated with a market-linked cluster.

We also found evidence supporting earlier emergence of a clade separated from lineage A by three mutations, one of which is found in closely related bat coronavirus genomes. Empirical analysis of SARS-CoV-2 phylogenetic trees showed that a similarly divergent clade is an unlikely, but plausible, coincidence. A divergent clade carrying a potentially ancestral mutation could emerge from a spillover from an unsampled population in a proximal host. However, Bayesian phylodynamic analysis using a multi-type birth-death model indicated that the ancestral mutation in this clade is more likely to be derived from lineage A than present in the sequence of the proximal SARS-CoV-2 ancestor. Remaining uncertainty might be resolved through the acquisition, publication, and analysis of additional sequences from early-pandemic samples, painting a clearer picture of SARS-CoV-2 emergence.

## Results

### Additional early-pandemic sequences do not increase support for alternative ancestral sequences or a single introduction

To construct our dataset, we started with a recently reported dataset of 863 sequences corresponding to samples collected between 24-Dec-2019 and 15-Feb-2020 [14]. In it, we identified 41 sequences to remove: 22 based on low reported coverage [21] and 19 identified as duplicates based on matching sample metadata in public databases and corresponding papers [22–25]. We added 187 sequences from public databases and new assemblies from public sequencing reads. We identified published studies corresponding to many of the added sequences [26–31]. Some sequences from Wuhan were published at the same time and in the same BioProject as sequences described in a genomic epidemiological study of Anyang [20]. Additional sequences from Beijing were added after deduplication based upon sample metadata in papers with overlapping groups of patients [32–36]. Details on databases where data can be found and corresponding accession numbers are available in **Supplemental Data 1**.

The final dataset is composed of 1,009 sequences. **Figure 1** shows their temporal distribution compared to the initial dataset (not including duplicates and low coverage sequences). Newly added data follows a similar distribution as the initial dataset starting in late January 2020. However, additional sequences collected until mid-January substantially add to the number of early samples, especially the number of samples without either a known market link or a link to a familial cluster in Guangdong province [15]. The earliest added sequence, collected on 7-Jan-2020 (YS8011), has a reported market link [26].

**Figure 1.**
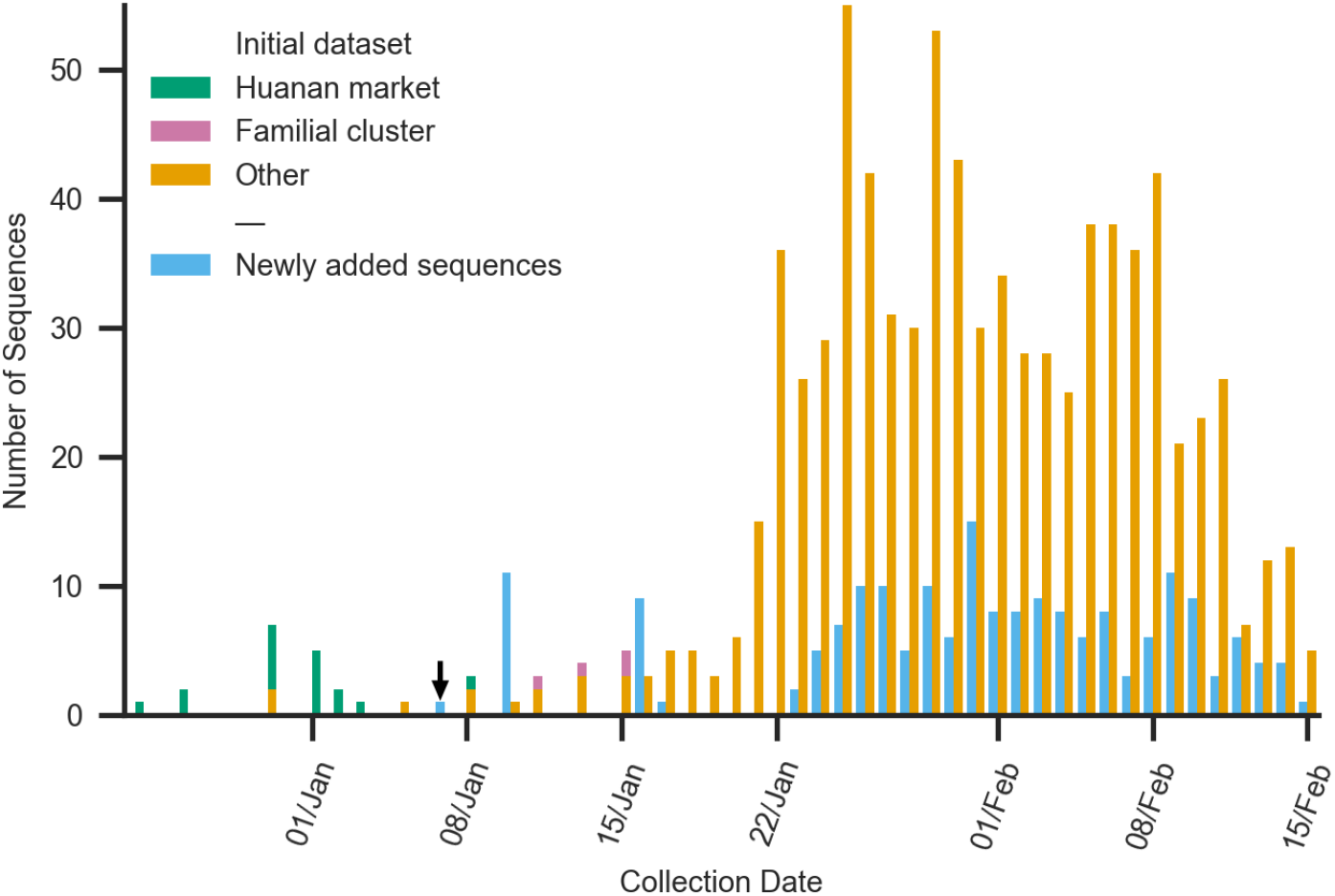
Temporal distribution of initial dataset and newly added sequences (samples collected between 24-Dec-2019 and 15-Feb-2020). Sequences in the initial dataset from market-linked samples and within a single familial cluster are colored green and purple, respectively. Other sequences in the initial dataset are colored orange. Newly added sequences are colored blue. The arrow indicates the earliest sequence in added data, which is market-linked.

For a first comparison of the composition of the initial and updated datasets, we compared the fractions of sequences in lineage A (sequences with mutations C8782T and T28144C compared to the SARS-CoV-2 reference sequence). This addresses the possibility that lineage A was under-ascertained in early-pandemic sequence data, since market-linked patient sequences were in lineage B. **Figure 2** shows that added data have an excess of lineage A sequences relative to the initial dataset starting in late January. However, this difference is driven by intensive sequencing in Anyang to characterize transmission chains [20]. The frequencies of lineage A sequences are plainly indistinguishable after excluding Anyang-focused sampling (52 sequences collected between 24-Jan and 14-Feb-2020), suggesting that additional data has a similar overall composition to the initial dataset.

**Figure 2.**
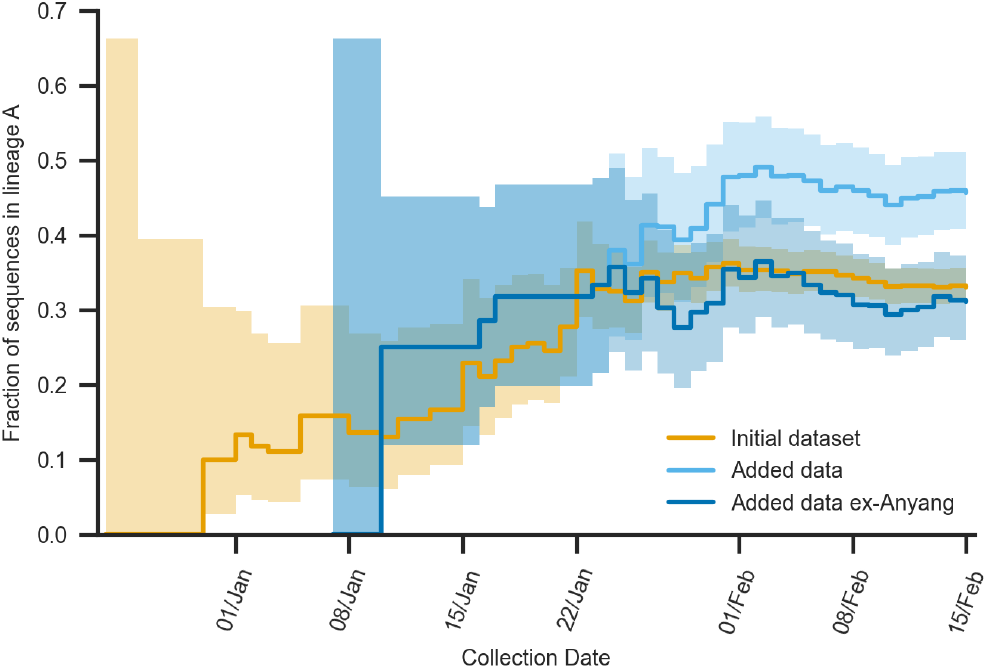
The cumulative fraction of sequences in lineage A for the initial dataset (orange), sequences added in this study (light blue), and added sequences excluding those collected in Anyang (dark blue). The blue lines are identical prior to the earliest sample from Anyang on 25-Jan-2020. With the exception of sequences from Anyang, newly added sequences have a similar composition of lineage A and B to sequences considered in previous analyses. Time series of binomial proportions with Wilson score intervals (z = 1.4) are shown as confidence bands.

In order to see whether additional data identifies any SARS-CoV-2 lineages as underrepresented in the initial dataset, we examined the distribution of newly added sequences in the context of a phylogenetic reconstruction of all 1,009 sequences (**Figure 3A**). Added sequences are distributed throughout lineage A and lineage B. We focused on lineage A sequences since the most recent common ancestor (MRCA) of the pandemic is likely in lineage A [10,13]. Two groups of added sequences in lineage A in late January and early February consist of samples from clusters in Anyang [20].

**Figure 3.**
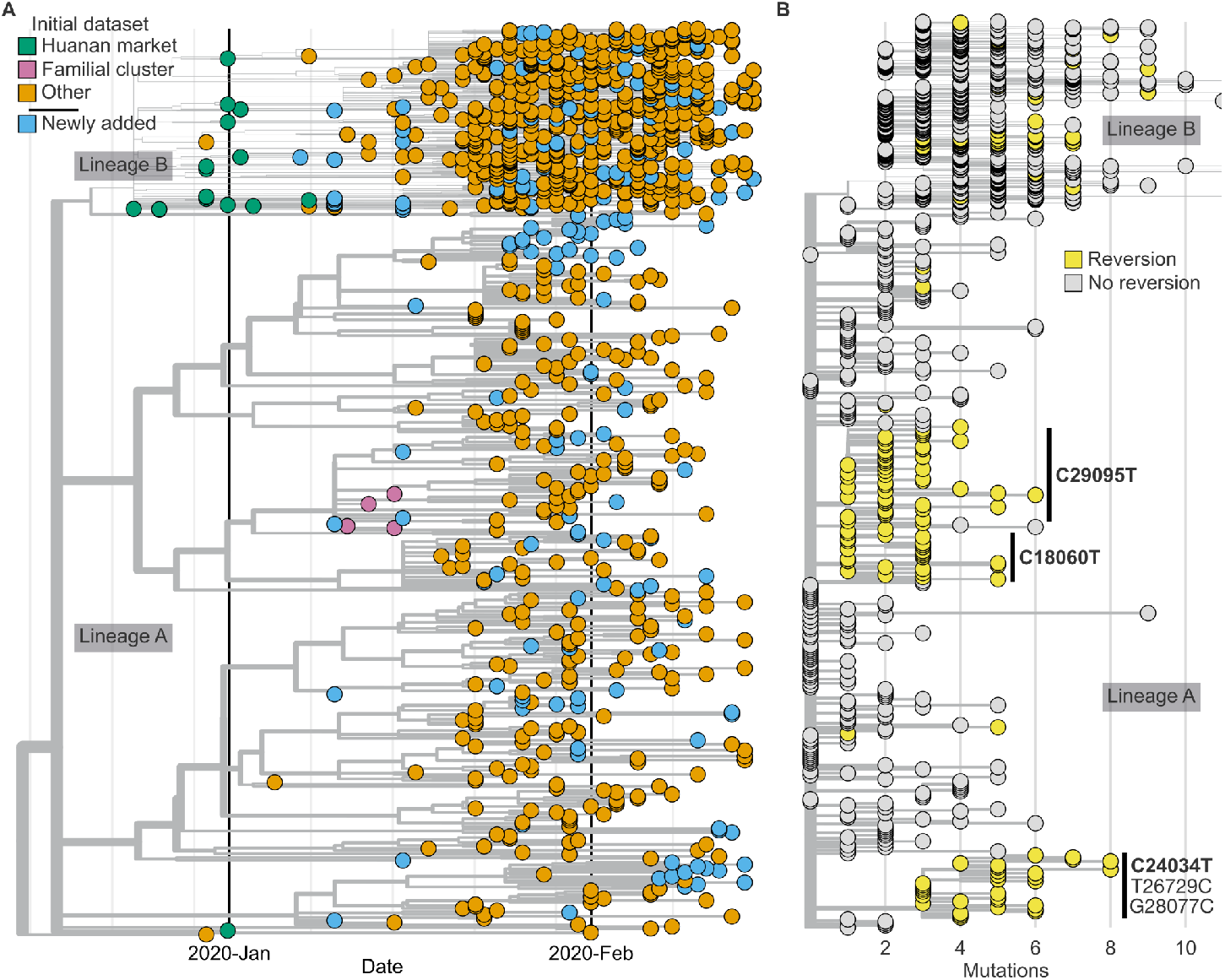
Phylogenetic trees of 1,009 samples collected between 24-Dec-2019 and 15-Feb-2020 are shown, rooted on lineage A. Lineage B samples, at the top of the trees, are vertically compressed to focus on lineage A. A given sequence is shown at the same vertical location on both panels. (**A**) Samples are plotted by time and colored by category: market-linked (green), linked to an early familial cluster (purple), other samples in the initial dataset (orange), and newly added samples (blue). (**B**) Samples are plotted by divergence from lineage A (a few lineage B samples over 10 mutations divergent from lineage A are not shown). Samples harboring mutations towards either the inferred ancestral sequence RecCA or C24034T are colored yellow. Three clades directly descending from lineage A are annotated; they contain 49 (C29095T), 24 (C18060T), and 32 (C24034T, T26729C, G28077C) descendant samples.

We first looked for potential intermediate sequences between lineage A and B (separated by the mutations C8782T and T28144C relative to the reference sequence Wuhan-Hu-1). We identified one sample, Guangzhou/ID098, collected on 31-Jan-2020, carrying C8782T and additional mutations G5062T, A9707G, and C29303T. However, this is not a sample from a transitional lineage, since it shares the mutation C29303T with other sequences in lineage A, including some collected in Guangzhou. We identified another sample, Guangzhou/ID106, collected on 31-Jan-2020, carrying T28144C. However, it similarly is not a transitional sequence since it shares C10604T, A15647G, and G29868A with other sequences in lineage B. Thus, this analysis of additional data, like previous analyses, found that intermediate sequences are either artifactual or derived from lineage A or B [13,14].

Next, we noted that previously proposed ancestral sequences in lineage A with mutations C18060T [17] or C29095T [16] were found in newly added sequences. These mutations are found in RaTG13 [37] and other closely related bat coronaviruses. It is likely that the recent ancestors of SARS-CoV-2 prior to the first human infection harbored one of these or another similarly “ancestral” mutation. However, phylodynamics analyses so far have found that these genotypes were unlikely to represent the MRCA of SARS-CoV-2 [10,13].

We examined the updated dataset to see if it was likely to increase support for C29095T (10 new sequences; earliest 10-Jan-2020) or C18060T (7 new sequences; earliest 23-Jan-2020) as being ancestral. We also investigated an early pandemic clade that was previously highlighted as bearing a reversion, C24034T, to bat coronaviruses RaTG13, RpYN06, and RmYN02 [16]. Although there were 13 newly added samples in the C24034T clade (earliest 10-Jan-2020), 9 of these were sampled in Anyang and share a lineage-defining mutation, T490A [20]. **Figure 3B** shows that these mutations are unique in characterizing relatively large clades in lineage A with potential ancestral mutations. Other reversions to sequences in related animal coronavirus mutations are also shown on **Figure 3B**, annotated for reversions to the inferred ancestral sequence RecCA [13].

For each of these mutations, additional data did not significantly change the overall composition of our dataset (**Figure 4**). Thus, additional data reinforces previous analyses finding it unlikely that the first human SARS-CoV-2 infection had C18060T or C29095T [10,13]. The mutation C24034T was not in the inferred ancestral sequence RecCA used in those analyses. The C24034T clade stands out as bearing three mutations (**Figure 3B**), and in the updated dataset its earliest sampling date moves from 15-Jan to 10-Jan–2020. The possibility that this divergent clade emerged from a spillover from an unsampled proximal host is discussed below.

**Figure 4.**
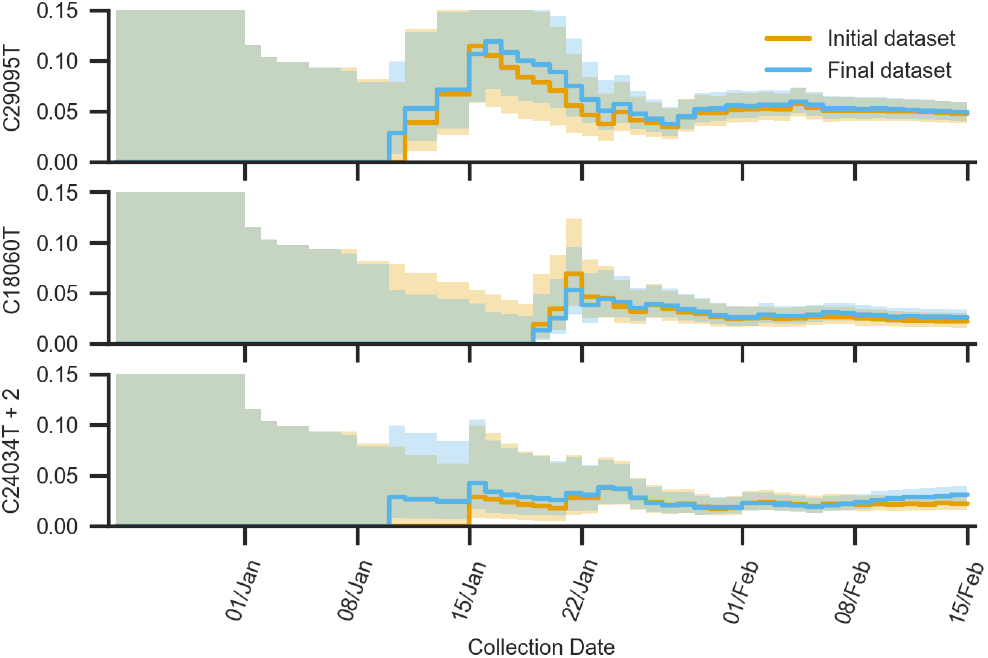
The cumulative fraction of the initial and updated sequencing datasets are shown for three clades with mutations from lineage A (C29095T, top, C18060T, middle, or all three of C24034T, T26729C, and G28077C, bottom). This shows how the overall relative composition of the dataset changes as samples are added over time. Time series of binomial proportions with Wilson score intervals (z = 1.4) are shown as confidence bands.

### Two early pandemic sublineages may have spread from the Huanan market

Sequences from market-linked patients were collected starting in late December 2019, several weeks after primary SARS-CoV-2 infection(s) seeded the outbreak [13]. If the market played a significant role in the early spread of SARS-CoV-2, mutations in market-linked patient sequences could appear in unlinked samples if transmission chains leading to them came from the market. In contrast, it would be less likely to detect market-linked mutations in unlinked patients under the hypothesis of an earlier origin elsewhere, according to which apparent market centrality reflects ascertainment bias [16]. To test this, we investigated the updated dataset for evidence of spread of mutations first identified in market-linked patients.

Mutations in common between market-linked and market-unlinked patients could signify transmission chains from the Huanan market. The report of the 2021 China-WHO joint mission in Wuhan [18] includes as Table 6 a list of mutations found in SARS-CoV-2 sequences from patients with Dec-2019 onset dates and a market link (they are all in lineage B sequences). Some initially identified mutations, still present in published consensus sequences (and included in our dataset), were not confirmed after reanalysis described in the WHO report. In our dataset, mutations, C6968A, T11764A, T13270C, and A24325G are found in sequences from market-linked patients with Dec-2019 onset. However, the WHO report found that C6968A and T11764A in sample WH01 could not be confirmed in reanalysis. In our dataset, T13270C only appears elsewhere in a clade unrelated to the market-linked sequence and sharing additional mutations C3885A, C12778T, and G29449T. Another market-linked mutation that could not be confirmed by reanalysis in the WHO report and is not in any market-linked sequence in our dataset, C28253T, is found in six sequences, but three of these sequences were clearly derived from other lineages without any market association, reflecting C28253T homoplasy. Since C28253T was identified as an intrahost variant in a market-linked sample [38], we concluded that it was unlikely that C28253T sequences in our dataset reflect spread from the Huanan market.

In contrast, the market-linked mutation A24325G appears eight times in our dataset, including five times in newly added sequences. Furthermore, the A24325G mutation was the only mutation we could identify shared by a cluster of market-linked patients in December 2019 (**Methods**), so we investigated whether additional sequencing data supported its spread from the market. Most previous analyses considered datasets in which there was only one A24325G sequence collected prior to 15-Feb-2020 with a market link: a sequence collected in California, USA on 12-Feb-2020 (CA-CDC-8). The recent publication of sequencing from Shanghai [19] added another collected on 6-Feb-2020 (SH-P243-2-Shanghai), and A24325G was also detected in a partial sequence collected in Wuhan on 30-Jan-2020 [24,39]. Newly added data considered in this study includes two A24325G sequences on 10-Jan-2020 and three others starting in late January. The earliest full-length sequence with A24325G without a known market link is now 27 days earlier, increasing the likelihood that these sequences reflect spread of this sublineage from the market.

Another market-linked patient was not included in the WHO report Table 6 because symptom onset occurred after Dec-2020. This patient visited clinics for treatment in Wuhan between 1-Jan and 4-Jan-2020, but was only later diagnosed with COVID-19 after returning to his hometown of Jingzhou [40]. The patient’s sequence (HBCDC-HB-01/2020) includes the C26370T mutation [41], which we found in one newly added sequence collected on 10-Jan-2020 in Wuhan and four sequences collected in China and Thailand starting in late January. Also of note, given the absence of reported market-linked patient sequences in lineage A, this study reported that another sequence in lineage A (HBCDC-HB-03/2020) came from a patient residing about 2 km from Huanan market, making it similar to two other early sequences in lineage A [2]. This is important because it adds additional evidence that early transmission of lineage A occurred near the Huanan market [5]. Together, newly added sequences in our dataset increase the likelihood that SARS-CoV-2 lineages carrying A24325G or C26370T spread from the market, further supporting the Huanan market as the pandemic epicenter.

### A divergent early-pandemic sublineage carries a potentially ancestral mutation

We asked whether new data would support any additional, distinguishable introductions of SARS-CoV-2 (i.e. with sequences other than lineage A or lineage B). As noted above, a previous analysis had annotated an early pandemic lineage harboring one mutation, C24034T, towards the bat coronavirus RaTG13 as well as two other mutations, with no transitional samples [16]. Multiple spillovers need not have high sequence diversity—for instance, the viral genomes RshSTT182 and RshSTT200 were collected from two different bats and differ by only 3 mutations [42]. However, branches with an unusual degree of divergence could arise from additional introductions from an unsampled reservoir with some genomic diversity. Although the presence of a mutation found in the most closely related bat coronaviruses could indicate an ancestral sequence, such a reversion could also occur by chance, as has occurred throughout the pandemic [13].

**Figure 4**. shows that the earliest sample in the C24034T clade was collected on 10-Jan-2020, the same day as the earliest sample for the previously proposed progenitor C29095T and nine days earlier than the earliest sample for C18060T. This is five days earlier than the earliest C24034T sample in our initial dataset, supporting earlier emergence of this clade. Phylogenetic reconstruction of the dataset of 1,009 samples revealed no evidence of samples with transitional sequences (lacking any of the three mutations characterizing this clade: C24034T, T26729C, or G28077C). Sample Hong_Kong/VM20001061-2 has ambiguous C24034Y (C or T) and shares C1663T and G22661T with another sequence from Hong Kong. Sample CHN/AY508 lacks G28077C, but shares T490A with 10 other sequences from China, South Korea, and the USA. Additionally, the C24034T clade was found in two samples in Wuhan in partial sequences from late January [39]. Thus, the only two samples lacking all three mutations were clearly not transitional sequences. This identifies the C24034T clade as a potential introduction carrying an ancestral mutation, which we now test.

**Figure 5**. shows a phylogenetic reconstruction of closely related human, bat, and pangolin coronaviruses in the region surrounding the C24034T mutation. Although C24034T was not included in the inferred ancestral sequence RecCA in previous work [13], it subsequently was found in closely related bat coronaviruses [43,44] and it now appears likely that a recent ancestor of SARS-CoV-2 carried C24034T. However, it is challenging to infer the recombinant history of this region of the genome; position 24034 falls in a variable and frequently recombinant genomic region within the sequence coding for the spike fusion peptide, following the S2′ proteolytic cleavage site.

**Figure 5.**
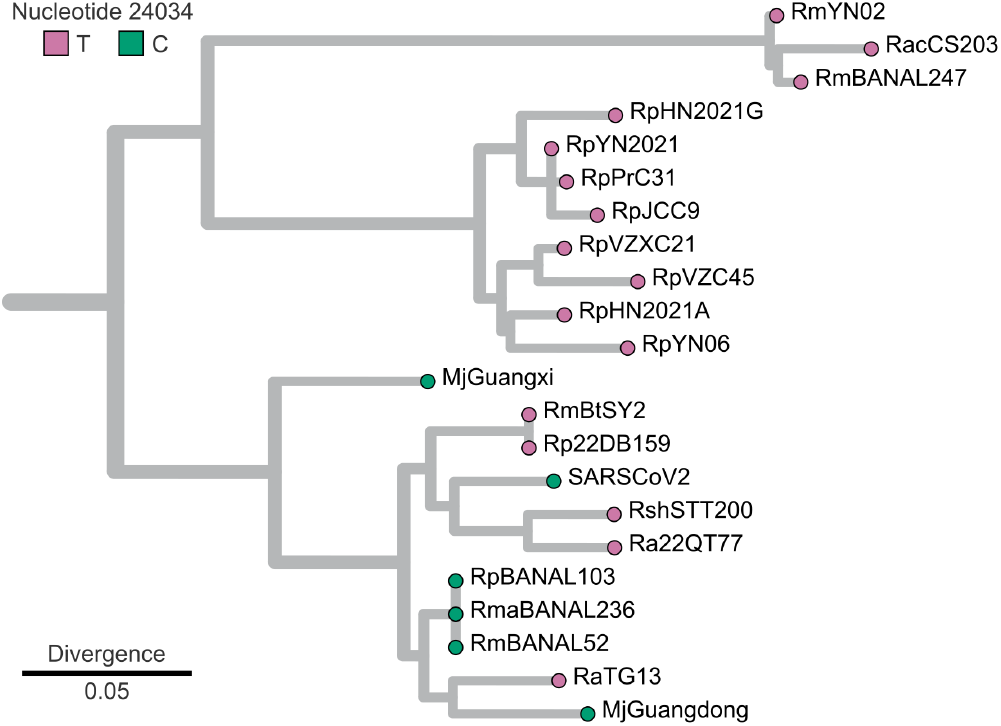
Phylogenetic tree of the region containing SARS-CoV-2 position 24034 and shown to have the most robust phylogenetic signal in previous analysis, and adding the sequence RmBtSY2, also called CX1 [43], to 111 human and animal sarbecovirus sequences considered in that study [44]. This corresponds to positions 23734–24183 in the SARS-CoV-2 reference genome. Only the sequences most similar to SARS-CoV-2 are shown. Samples are colored by nucleotide identity at the equivalent position to SARS-CoV-2 24034 in the multiple sequence alignment.

Although the C24034T clade stands out as a divergent clade within the early-pandemic phylogenetic tree (**Figure 3B**), we first empirically characterized the rarity of a similar topology in SARS-CoV-2 phylogenetic trees. We noted that there were three potential reversions (C18060T, C24034T, and C29095T) that gave rise to small clades, and we estimated the likelihood that one of three would form a similarly divergent polytomy by chance in the GISAID Global Phylogeny (12.6 million sequences). We found that 1.83% of clades similarly sized or larger (4,728 of 169,599) were separated by three or more mutations from their parent node. Accounting for the fact that any of the three common early-pandemic reversions could have been in a divergent clade by chance, this results in a 5.49% chance that one clade out of three would be this divergent from lineage A.

A similar analysis can be applied to the early-pandemic split between lineages A and B, which were separated by two mutations and formed large, similarly sized polytomies. This tree topology appeared in 3.1% of simulated epidemics [13]. In the GISAID phylogeny, 2.99% of polytomies with at least 20 children had a similarly sized child node (size ratio no more than three) with two additional mutations (2,626 of 87,691). These empirical analyses confirm the rarity of these features of the SARS-CoV-2 phylogenetic tree under the hypothesis of a single spillover; they motivate considering an alternate model of multiple spillovers from a diverse proximal reservoir.

### Phylodynamic analysis does not support any alternative ancestral genotype

Having shown that the divergence of the C24034T clade from lineage A was unlikely to happen by chance in the observed human SARS-CoV-2 tree, we next investigated the hypothesis that this clade was separately introduced from an unsampled proximal host. We also aimed to compare the relative likelihoods of the proposed ancestral sequences. We inferred parameters and phylogenetic structure in a multi-type birth-death model recently applied to MERS sequencing data [45]. In this model, infections propagate through competing processes of reproduction (infection of one host by another of the same type, or by a different type in spillover events) and removal (end of infectious period). We aimed for model simplicity given phylogenetic uncertainty, examining sequences collected prior to 23-Jan-2020, a period in which the sample size exhibits exponential growth (**Methods**).

We posited the existence of a sample from the proximal reservoir of human SARS-CoV-2 infections with a date of 15-Nov-2019 (a plausible date of primary human infection(s) [13]) and a lineage A sequence with unknown nucleotides (N) at positions 18060, 24034, and 29095. This choice essentially neglected scenarios in which the pandemic MRCA is lineage B or one of two possible transitional sequences, because we focused on the relative likelihoods of lineage A introduction or one of the three genotypes with one of C18060T, C24034T, or C29095T. This model is agnostic to the nature of the proximal reservoir other than being approximated by a birth-death process and plausibly being sampled on 15-Nov-2019. Exploration of prior distributions indicated that such a model was sensitive to assumptions about spillover rates (**Figures S1** and **S2**), so caution is required when considering absolute likelihoods that particular clades originated from spillover events. Furthermore, our site model did not account for irreversible substitution rates, an important consideration for SARS-CoV-2 phylogenetic inference [13]. However, since all substitutions of interest were C→T, we were able to compare the relative support for different ancestral sequences.

We found that Lineage B was monophyletic in 99.3% of sampled trees (N=90,001 sampled trees after discarding burn-in), and that it was introduced via one or more spillover events in 69.1% of those trees (overall, 82.6% of trees had multiple spillovers including trees with multiple spillovers in lineage A, but no lineage B spillover). Although this is consistent with a previous analysis concluding that lineage B and lineage A likely emerged from multiple introductions [13], it will be important to evaluate the robustness of this result in future work. Our analysis identified potential multiple spillovers by inferring the host type of internal nodes. The highest independent posterior subtree reconstruction (HIPSTR) supported two introductions, one for lineage A and one for lineage B **(Figure 6A**).

**Figure 6.**
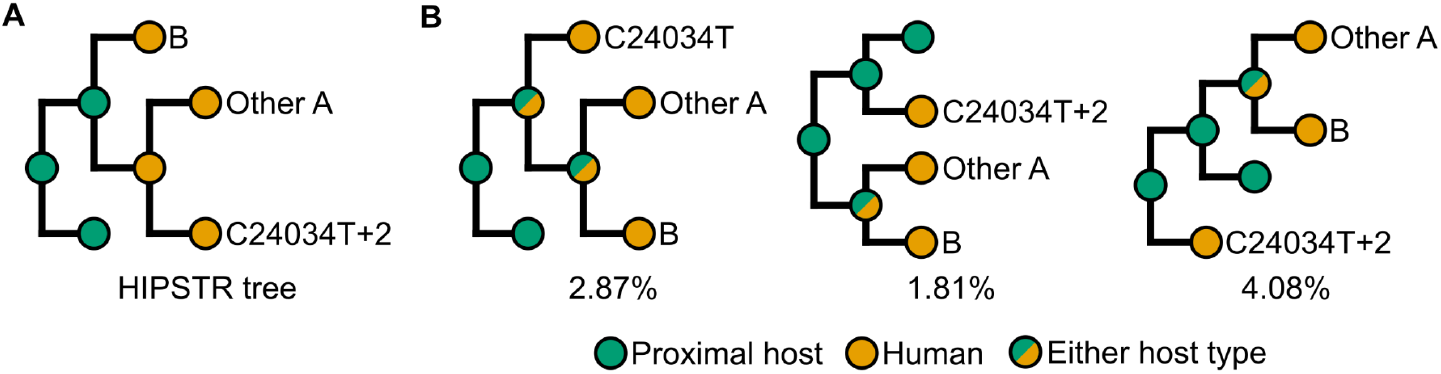
Consensus tree topology and alternative tree topologies in phylodynamics analysis. (**A**) Topology and internal node types of the highest independent posterior subtree reconstruction (HIPSTR) tree. Nodes colored by most likely type (proximal host, green, or human, orange). In this topology, the C24034T clade is sister to other lineage A sequences, all lineage A sequences are monophyletic, and the synthetic proximal host sample (terminal green node) is an outgroup to human samples. (**B**) Three topologies in which the C24034T clade is evolutionarily distinct from the MRCA of other lineage A sequences and lineage B (i.e. an outgroup to the clade containing lineage B and remaining lineage A sequences). Nodes are colored by possible types; the second two topologies have at least two spillovers. Some nodes are shown with split coloring to illustrate how the same topology can have different numbers of spillovers. Percentages indicate the frequencies of these topologies in the posterior tree distribution (N=90,001 sampled trees after discarding burn-in).

We compared the plausibility of proposed ancestral sequences in single- or multiple-spillover scenarios. We measured the likelihood in the posterior tree distribution that a clade of interest (C18060T, C24034T, or C29095T) was evolutionarily distinct from the MRCA of all other sequences (i.e. an outgroup to the clade containing lineage B and remaining lineage A sequences, as illustrated in **Figure 6B**). Since this does not require that the ancestral state includes this mutation, we also included a clade defined by the mutation C15480T as an estimate of how frequently these topologies occur by chance; C15480T was chosen as a well-supported clade (N=4 sequences) with a single C→T mutation in lineage A. Results are summarized in **Table 1**. Despite the limitations of very different methods in this work, results were comparable to an analysis constrained to RecCA (which lacks C24034T) that found 0.7% and 0.3% likelihoods for C29095T and C18060T pandemic MRCA sequences, respectively [10]. That analysis also found low, but non-zero likelihoods for T26729C or G28077C MRCA sequences (the two mutations occurring together with C24034T). This showed that an unusually divergent lineage modestly increased support for an ancestral genotype even without an ancestral mutation.

**Table 1.** Fraction of posterior tree distribution (N=90,001 after discarding burn-in) with a tree topology in which a specified clade is evolutionarily distinct from the MRCA of all other sequences; the total outgroup likelihood is sum of likelihoods of the three topologies shown in **Figure 6B** and can occur in trees with multiple or single spillovers. The clade spillover likelihood is the likelihood that the parent node of each clade is the proximal host type regardless of the tree topology. The C24034T clade additionally has mutations T26729C and G28077C.

|  | Clade |  |  |  |
| --- | --- | --- | --- | --- |
|  | C24034 + 2 | C29095T | C18060T | C15480T |
| **Total outgroup likelihood** | **8.75%** | **1.63%** | **0.74%** | **0.47%** |
| Multiple spillover | 7.72% | 1.35% | 0.63% | 0.40% |
| Single spillover | 1.04% | 0.28% | 0.11% | 0.07% |
| **Clade spillover likelihood** | **38.07%** | **20.90%** | **14.64%** | **13.06%** |

Previous work discussed the likelihoods of candidate progenitors in single-introduction scenarios, focusing on the mutations C18060T and C29095T [16,17]. However, we found that trees with topologies consistent with such alternative ancestry were no less likely to have multiple introductions than other trees (**Table 1**). The clade with C24034T was much more likely to be an outgroup to other sequences than clades with C18060T or C29095T, showing that it is a more plausible SARS-CoV-2 progenitor genotype. However, it was most likely that none of these three mutations characterized the progenitor. Furthermore, since our model did not account for the high rate of C→T mutations relative to T→C mutations, outgroup and spillover likelihoods in **Table 1** should be conservatively considered as upper limits. Thus, it is plausible, but unlikely, that the C24034T clade was an additional spillover, or that the MRCA of human SARS-CoV-2 had C24034T. The most likely scenario was that all clades were derived from lineage A.

## Discussion

We assembled an updated dataset of early-pandemic SARS-CoV-2 sequences, identifying sequences for exclusion and adding additional sequences not considered in previous analyses. This provided an opportunity to see whether or not previous conclusions were robust to adding a substantial amount of data—approximately doubling the number of sequences collected by mid-January 2020 without a reported link to the Huanan market or an early familial cluster. We found no intermediate sequences between lineages A and B and no significant change in the relative composition of these two lineages. Finding either, especially in early samples, would reduce the likelihood that there were two or more introductions of SARS-CoV-2, but this was not the case. We found no indication that additional data significantly changed the fraction of C29095T or C18060T in our dataset. Finding an increased frequency for either would have increased the likelihood of an alternative ancestral sequence, but this was not the case.

Rather than overturning previous results, newly added sequences and annotation of sample metadata made it possible to identify increased support for two sublineages of lineage B spreading from the Huanan market. This would be unlikely to occur if the market outbreak started as a small part of a wider outbreak. This is in addition to the low likelihoods that, by mere coincidence, both lineage A and lineage B would be found in market environmental sampling, and that three of the earliest lineage A sequences would be from patients located near the market. Conversely, all of these observations are straightforward predictions from the hypothesis that SARS-CoV-2 emerged from the Huanan market wildlife trade, a hypothesis first posed at the end of 2019 on social media before almost any data existed to test it [3,4].

We also identified a clade separated by three mutations from lineage A including one potentially ancestral mutation. Multi-type phylodynamic analysis suggested that this mutation characterizes a plausible, but unlikely, proximal ancestor genome. This part of our analysis could be improved by incorporating realistic spatiotemporal heterogeneities in transmission and sampling likelihoods as well as a more realistic substitution model, possibly utilizing empirical estimates of site-specific substitution rates [46]. Although our model was agnostic about the nature of the proximal host, a market-specific model could additionally incorporate a location trait for market-linked sequences.

More complex analyses would benefit from more complete sample metadata, especially with respect to market exposure and focused sequencing of patient clusters. An important limitation in this work is that sequences without a known market link may, in fact, have unpublished, direct links to the Huanan market. Epidemiological metadata was key, for example, to attribute some features of our dataset to focused sampling in Anyang [20]. The dataset reported here can also certainly be improved with additional scrutiny. For example, we have not removed duplicates from two studies that collected and sequenced independent samples from overlapping sets of patients [19,47]; amending published metadata to include cross-referenced patient ID numbers would make make it possible to construct a more accurate dataset. Additionally, analysis of early pandemic genomic diversity will be improved by publishing additional sequences such as those from samples collected in January 2020 [7].

Our analysis demonstrates how evidence continues to accumulate and clarify the origin of SARS-CoV-2. This is not limited to human SARS-CoV-2 sequences, but also to sequences of animal viruses such as those supporting C24034T as a reversion to an ancestral sequence. In addition to viral genome sequencing data, we note that other sources of data are often overlooked in commentary questioning the strength of the evidence for SARS-CoV-2 emergence in the Huanan market. These other datasets include spatiotemporal data of healthcare worker infections showing spread from the area around the market [6] and serosurveys of blood donor samples failing to find a broader early outbreak [48,49]. A combination of additional data and models that can synthesize diverse sources of data will continue to refine our understanding of SARS-CoV-2 origin.

## Methods

### Curation of sequence dataset

Our initial dataset was a recently reported dataset of 863 genomes collected between 24-Dec-2019 and 15-Feb-2020 [14]; we gathered assemblies corresponding to sample names in analysis configuration files available at https://github.com/pekarj/SC2_intermediates. To identify some additional sequences, we searched the GISAID database for newly published sequences and searched the RCoV19 resource (https://ngdc.cncb.ac.cn/ncov/) for sequences not in GISAID or GenBank databases. Other published assemblies were identified from publications as detailed in the **Results** section. **Supplementary Data 1** includes information for the database, assembly accession, related BioProject accession (where available), corresponding manuscript (where one was identified), and sample collection date. Collection dates were obtained from linked BioSample metadata. Mid-January 2020 sequences from Wuhan were published in January 2024 to BioProject PRJCA002163 together with sequences from Anyang [20]; BioSample data had been published one year earlier.

New assemblies in the dataset were constructed from sequencing reads associated with one or more of several related publications on samples from Beijing patients [32–36], some of which were made public at our request. Samples were deduplicated and collection dates were identified by cross-referencing experiment and BioSample metadata with supplementary tables in related manuscripts, retaining the earliest high-quality sequence for each patient. We assembled sequences from all samples from several related BioProjects and our dataset includes sequences from CRA002626 (15), HRA000181 (26), HRA000349 (15), and PRJNA667180 (1). Unmapped reads were mapped with Minimap2 [50] (options: -ax sr). Consensus sequences were assembled using ViralConsensus [51] (options: -q 20 -d 10 -f 0.5). Assemblies were subsequently refined with a 75% consensus threshold. Sequences with less than 95% coverage were excluded from the dataset for consistency with methods used to construct the initial dataset [13].

Additionally, some samples in the initial dataset were removed. **Supplementary Data 1** describes the rationale for removal such as evidence supporting sequences as duplicates. Other samples were excluded because of coverage below 95% according to supplementary data in a related publication [21]. We also made a modification for which market-linked sequences were included in our dataset. Although A24325G was reportedly found in consensus genomes for two market-linked patients (S04 and S12 in Table 6 of the Joint WHO-China Study [18]), 1 of 5 sequences for S04 lacks

A24325G (IME-WH02), and only 1 of 2 sequences for S12 has A24325G (IME-WH03). We concluded that swapping IME-WH02 and IME-WH03 was most parsimonious with available sequence data and metadata; a similar change was previously made for other sequence-patient pairs in the report [52]. For patient S12, we opted for IME-WH02 over WIV06 in our dataset due to higher reported sequencing depth in the Joint WHO-China Study [18]; both sequences contain no substitutions relative to lineage B. This left the sequence for patient S04 (WH19008) as the only market-linked sequence with A24325G in our dataset. Further inspection of sequencing and metadata [38] identified another sequence, WH19003, containing A24325G and likely corresponding to a different patient in the same cluster as WH19008 (Cluster 2 in the Joint WHO-China Study). Although coverage is too low to include sample WH19003 in our dataset, together this identifies A24325G as the only mutation shared by a cluster of patients in Huanan market in late December 2019.

### Phylogenetic analyses

The entire dataset of 1,009 sequences was analyzed using the Nextstrain ncov pipeline [53] available at https://github.com/nextstrain/ncov, rooting on lineage A to infer the phylogenetic tree and date samples collected between 24-Dec-2019 and 15-Feb-2020. Annotation was added based upon mutations in RecCA [13] identified by Nextclade [54], with the addition of C24034T. Auspice was used to visualize trees for both **Figure 3** and **Figure 5**.

To investigate the C24034T mutation, we first used MAFFT [55] to add BtSY2, also known as CX1 [43], to a multiple sequence alignment of 111 animal and human coronaviruses [44]. The region of the alignment corresponding to the phylogenetic coloured genomic bootstrap (CGB) barcode around SARS-CoV-2 position 24034 was extracted from the alignment. IQTree [56] (options: -st DNA -m GTR+F -czb --keep-ident -nt 4 -redo -seed 2 -asr) was used to construct the phylogenetic tree of this region, and only a portion of the tree is shown in **Figure 5** for clarity.

We analyzed SARS-CoV-2 phylogenetic trees with Python scripts using the ETE Toolkit [57] to estimate the likelihood that one of three potentially ancestral mutations would form a similar divergent polytomy. Each node we examined had at least 26 descendants, matching the number of C18060T samples in our dataset. In the GISAID Global Phylogeny [58], 1.83% of these nodes formed polytomies at least three mutations diverged from their parent with 10 or more children. This was similar to 1.81% in the public UShER tree [59] and 2.69% for an early-pandemic phylogenetic tree [60]. Little change in likelihood was observed as a function of distance from the root of the tree. The likelihood that two similarly sized polytomies would be separated by 2 mutations was analyzed similarly according to the polytomy size criteria described in **Results**.

Bayesian phylodynamics analysis utilized the BDMM-Prime multi-type birth-death model [45,61,62] and BEAST 2.5 [63]. We used sequences collected on or before 22-Jan-2020 (N=129; sample YB20200116082 was excluded based upon having 9 substitutions between positions 15101 and 15111). The cutoff date was chosen based upon approximately exponential growth of the cumulative number of sequences through this date, allowing for a relatively simple model of sampling likelihood while including several samples (N>=5) for each of the C18060T, C24034T, and C29095T clades. Sequences were aligned using NextClade [54]. Positions 1–88 and 29709–29903 were masked based upon lacking coverage in at least 5% of samples. A synthetic sample dated 15-Nov-2019, having N at positions 18060, 24034, and 29095, and otherwise identical to lineage A sample WH04 was added, positing that a sample with this ambiguous haplotype could have been collected at this time from the proximal reservoir of the human SARS-CoV-2 outbreak. This was a plausible date of primary human SARS-CoV-2 infections in previous analysis [13]. Model specification is described in **Supplementary Data 2**. To simplify the model given phylogenetic uncertainty, we constrained some parameters e.g. using a fixed death rate as in previous work [64].

Posterior tree distributions were analyzed to construct HIPSTR reconstructions using TreeAnnotator [65]. Topologies and host-type likelihoods for parent nodes of clades of interest were quantified with Python scripts utilizing TreeSwift [66]. We discarded 10% of 100,001 trees (sampled every 1,000 of 100 million iterations) and confirmed effective sample sizes above 200 for all inferred parameters using Tracer [67]. **Figures S1** and **S2** show results of a sensitivity analysis in which we repeated analysis using different prior distributions for the spillover rate, parameterized as an effective reproduction number. Absolute likelihoods that clades were founded by spillovers or had ancestral topologies varied somewhat, but relative likelihoods consistently found C24034T much more likely than C18060T or C29095T to be ancestral.

## Data Availability

Our sequencing dataset cannot be published, since it contains sequences exclusively available from GISAID. We provide as **Supplementary Data 1** a spreadsheet with information required to reproduce our sequencing dataset. Likewise, phylodynamics configuration files contain sequence data from GISAID; modeling parameters required to reproduce our results are available in **Supplementary Data 2**.

## Acknowledgments

We thank Hui Zeng and colleagues at Beijing Ditan Hospital, Capital Medical University for making available sequence data in GSA for Human repositories HRA000181 and HRA000349. They are gratefully acknowledged along with all other data contributors, i.e., the authors and their originating laboratories responsible for obtaining the specimens, and their submitting laboratories for generating the genetic sequence and metadata and sharing via GISAID and other databases, on which this research is based. **Supplementary Data 1** includes names of submitting laboratories and individuals. To view the contributors of each individual sequence for sequences used from GISAID, visit https://doi.org/10.55876/gis8.250331rs. We thank Niema Moshiri, Jonathan E. Pekar, and Alexander Crits-Christoph for advice on analysis and critical comments on this manuscript.

## Supplemental Figures

**Figure S1.**
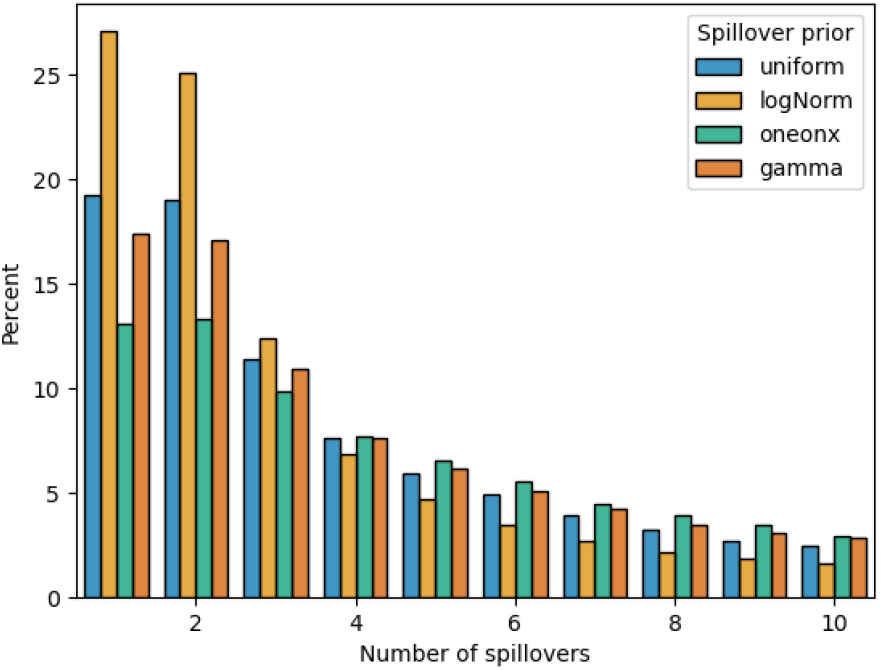
Histograms of the number of spillovers in the posterior tree distribution, colored by the prior distribution used for the effective reproduction number for spillover from the proximal host to humans. Trees are likely to have multiple spillovers with all prior distributions that were tested.

**Figure S2.**
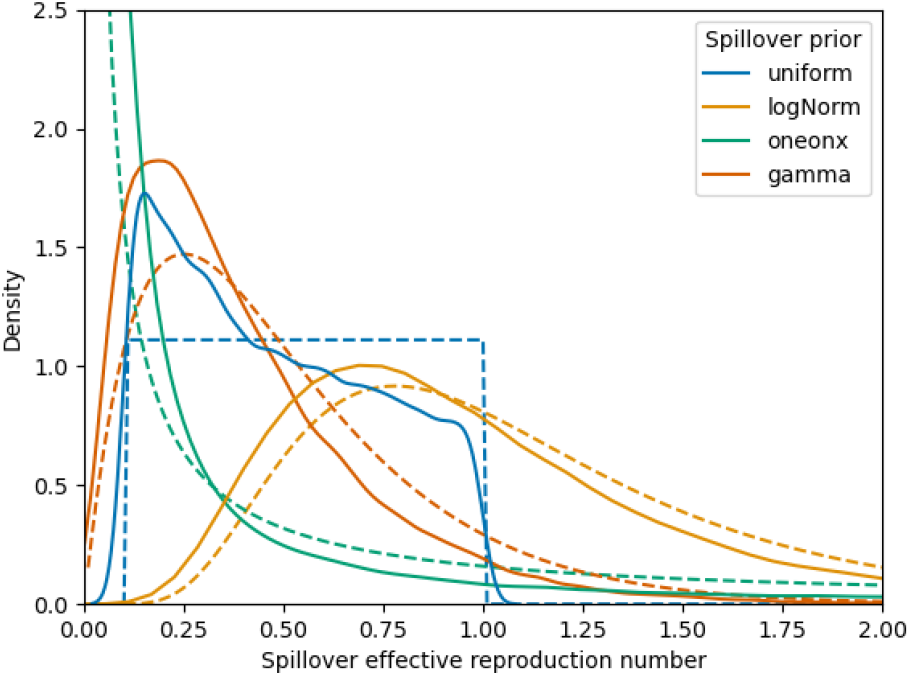
Prior distributions (dashed) and kernel density estimates of posterior distributions (solid) for the effective reproduction number for spillover from the proximal host to humans. Colored by the prior distribution used. Small differences between prior and posterior distributions indicate sensitivity to prior specification.

